# Classical swine fever virus NS5A inhibits NF-κB signaling by targeting NEMO

**DOI:** 10.1101/2021.06.21.449351

**Authors:** Jiaying Li, Haixiao Yu, Wenjin Jiang, Ping Ma, Zezhong Feng, Yang Lu, Changchun Tu, Jinfu Sun

## Abstract

The NS5A non-structural protein of classical swine fever virus (CSFV) is a multifunctional protein involved in viral genomic replication, protein translation and regulation of cellular signaling pathways, and assembly of infectious virus particles. Previous report showed that NS5A inhibited nuclear factor kappa B (NF-κB) signaling induced by poly(I:C); however, the mechanism was not elucidated. Here, we report that NS5A interacts with NF-κB essential modulator (NEMO),a regulatory subunit of the IκB kinase (IKK) complex, and that the zinc finger domain of NEMO essential for NEMO ubiquitination and IKK activation is required for the interaction of NEMO with NS5A. Viral infection or NS5A expression by itself reduced the protein level of NEMO. Mechanistic analysis revealed that NS5A mediated proteasomal degradation of NEMO. Ubiquitination assay showed that NS5A induced K27-but not K48-linked polyubiquitination of NEMO. In addition, NS5A blocked k63-linked polyubiquitination of NEMO, thereby inhibiting activation of IKK and NF-κB. These findings indicate that NS5A inhibits NF-κB signaling by mediating proteasomal degradation of NEMO and blocking k63-linked polyubiquitination of NEMO, thereby revealing a novel mechanism by which CSFV inhibits host innate immunity.

**Importance:** Classical swine fever (CSF) is a economically important swine viral disease leading to hemorrhagic fever and immuno-suppression. In order to successfully infect and replicate in a host cell, viruses have evolved various strategies to antagonize host innate immunity. It is known that CSFV non-structural protein N^pro^ interacts with interferon regulatory factor 3 (IRF3) and mediates its proteasomal degradation, thereby inhibiting the production of type I interferon. However, no other mechanism by which CSFV antagonizes host innate immunity has so far been reported. Here, we show that NS5A inhibits NF-κB signaling by mediating proteasomal degradation of NEMO and by blocking k63-linked polyubiquitination of NEMO, thereby revealing a novel mechanism by which CSFV antagonizes host innate immunity.

## Introduction

Classical swine fever (CSF) is a economically important swine viral disease characterized by acute hemorrhagic fever, high morbidity and mortality or atypical, inapparent clinical manifestations depending on the virulence of the virus strain (1). The causative agent, CSF virus (CSFV) is a member of the genus *Pestivirus* within the family *Flaviviridae* (2). The CSFV genome contains a large open reading frame (ORF) that encodes a polyprotein which is cleaved by cellular and viral proteases into 4 structural proteins, C, E^rns^, E1, and E2, and 8 nonstructural proteins, N^pro^, p7, NS2, NS3, NS4A, NS4B, NS5A and NS5B (3).

CSFV NS5A is a multifunctional protein involved in viral RNA replication, protein translation and virus assembly. NS5A regulates viral RNA replication by binding to the 3’-UTR and NS5B (4,5) and functions in the switch from translation to replication by interaction with NS5B (6,7). NS5A also participates in virus assembly by interaction with core protein (8) or with host proteins such as annexin A2 (9), Rab1A (10) and Rab18 (11). Our previous report indicated that NS5A interacts with host eIF3E, thereby facilitating viral RNA replication and protein translation, and inhibiting host translation (12). Cellular Hsp27 (13), eEF1A (14) and viperin (15) have been reported to interact with CSFV NS5A and negatively regulate viral replication. Additionally, two other NS5A binding partners, HSP70 (16) and FKBP8 (17), have been shown to facilitate viral proliferation. NS5A also inhibits the secretion of inflammatory cytokines induced by poly(I:C) by suppression of the NF-κB signaling pathway (18). However, the mechanism of inhibition of the NF-κB signaling by NS5A remains obscure.

Nuclear factor kappa B (NF-κB) plays a critical role in inflammatory immune responses, cell growth and apoptosis (19). In unstimulated cells, NF-κB is sequestered in the cytoplasm through its association with inhibitory IκB proteins which mask its nuclear localization signal. Following stimulation by cytokines such as tumor necrosis factor (TNF)-α or interleukin (IL)-1, IκB kinase (IKK) is activated and phosphorylates IκBs which are polyubiquitinated and further degraded by 26S proteasomes to allow NF-κB to translocate to the nucleus and bind to a specific sequence to activate target gene transcription. The IKK complex consists of two catalytic subunits, IKKα and IKKβ, and a regulatory subunit NF-κB essential modulator (NEMO or IKKγ) which contains two coiled-coil domains, CC1 and CC2, a leucine zipper (LZ), a ubiquitin-binding domain, and a C-terminal zinc finger (ZF), and plays a critical role in the activation of IKK and NF-κB (20-22).

Upon viral infection, cellular pattern recognition receptors (PRRs) sense viral pathogen-associated molecular patterns (PAMPs). This rapidly induces host innate and subsequent adaptive immune responses, leading to viral suppression (23). The activation of NF-κB plays a critical role in these early events and in the inflammatory responses following microbial infection (24-26). Viruses have therefore evolved various mechanisms targeting the NF-κB signaling pathway to evade the host innate immune response. Here, we show that CSFV NS5A inhibits the NF-κB signaling pathway by mediating proteasomal degradation of NEMO and by blocking K63-linked polyubiquitination of NEMO.

## Results

### NS5A interacts with cellular NEMO

In our previous study, screening for host partners binding to CSFV NS5A was performed using the His-tag pull-down assay and LC-MS/MS identification (12). For the present study, NEMO, one binding partner of NS5A, was selected for further analysis because of its critical roles in the NF-κB signaling pathway. To confirm the interaction of NS5A with NEMO, co-immunoprecipitation (Co-IP) analysis was conducted. 293T cells expressing FLAG-NS5A fusion protein (Flag tag; Fig.1 A) and 293T cells expressing His-NEMO fusion protein (His tag and FLAG-NS5A fusion protein; Fig.1 B) were lysed and immunoprecipitated with mouse anti-Flag monoclonal antibody (Proteintech, Wuhan, China) (Fig.1 A) or mouse anti-His tag monoclonal antibody (Abcam, USA) (Fig.1 B). Precipitated proteins and whole cell lysates were subjected to western blotting with rabbit anti-Flag polyclonal antibody (Proteintech, Wuhan, China) and rabbit anti-NEMO monoclonal antibody (Abcam, USA) (Fig.1 A) or rabbit anti-FLAG polyclonal antibody and mouse anti-His tag monoclonal antibody (Abcam, USA) (Fig.1 B) respectively. Results indicate that NEMO specifically co-precipitated with the fusion protein FLAG-NS5A but not Flag tag (Fig.1 A), and FLAG-NS5A co-precipitated with the fusion protein His-NEMO but not His tag (Fig.1 B).

**Fig. 1.**
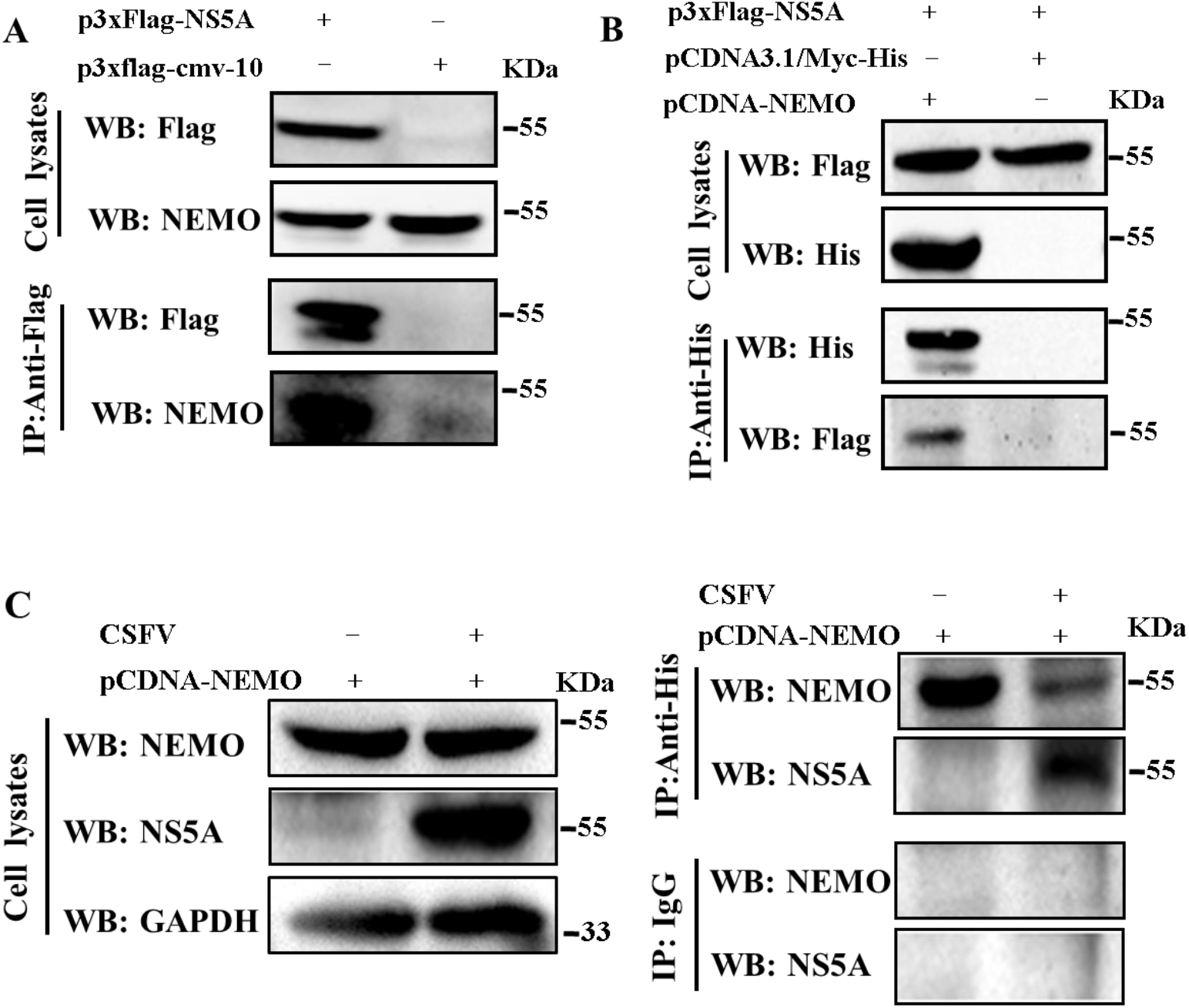
Evidence that CSFV NS5A interacts with NEMO. (A, B) 293T cells were transfected with indicated plasmids. After 48 h, cells were lysed for immunoprecipitation (IP) and western blot assay with indicated antibodys. (C) PK15 cells were infected with CSFV Shimen strain at MOI of 3 TCID50 and transfected with pCDNA-NEMO vector. At 48 hpi, IP and western blot assay were conducted using the indicated antibodies.

PK-15 cells infected with CSFV and transfected with pCDNA-NEMO were lysed and cell extracts were immunoprecipitated with mouse anti-His tag monoclonal antibody (Abcam, USA) or nonspecific IgG. Precipitated proteins and cell lysates were then subjected to western blotting with rabbit anti-NS5A serum (prepared in our laboratory) and rabbit anti-NEMO monoclonal antibody (Abcam, USA). Fig1C shows that the native NS5A produced by virus infection also co-precipitated with NEMO; i.e., both Flag-tagged and native NS5A interacted with NEMO.

### The domain of NEMO required for NEMO-NS5A interaction

To determine the region of NEMO required for interaction with NS5A, recombinant vectors expressing distinct deletion mutants of NEMO (Fig 2A) were constructed. His-NS5A fusion protein was expressed and purified as previously described (12). Separate cultures of 293 T cells were transfected with recombinant plasmid pCMV-NEMOΔN1, pCMV-NEMOΔN2, pCMV-NEMOΔN3 or pCMV-NEMOΔN4. At 48 h post-transfection, the cells were lysed and incubated with Dynabeads (Thermo Fisher Scientific, USA) coupled with His-NS5A or His tag. Bound proteins and cell lysates were then subjected to western blot analysis with anti-Myc antibody. Fig 2B shows that the NEMOΔN3 mutant interacted with NS5A, indicating that the 1-125aa N terminal region of containing the IKKα/β binding domain(res idues 44-86aa)(27) of NEMO was unnecessary for the interaction between NS5A and NEMO; however, regions 126-220aa and 351-419aa containing the ZF domain were at least required.

**Fig. 2.**
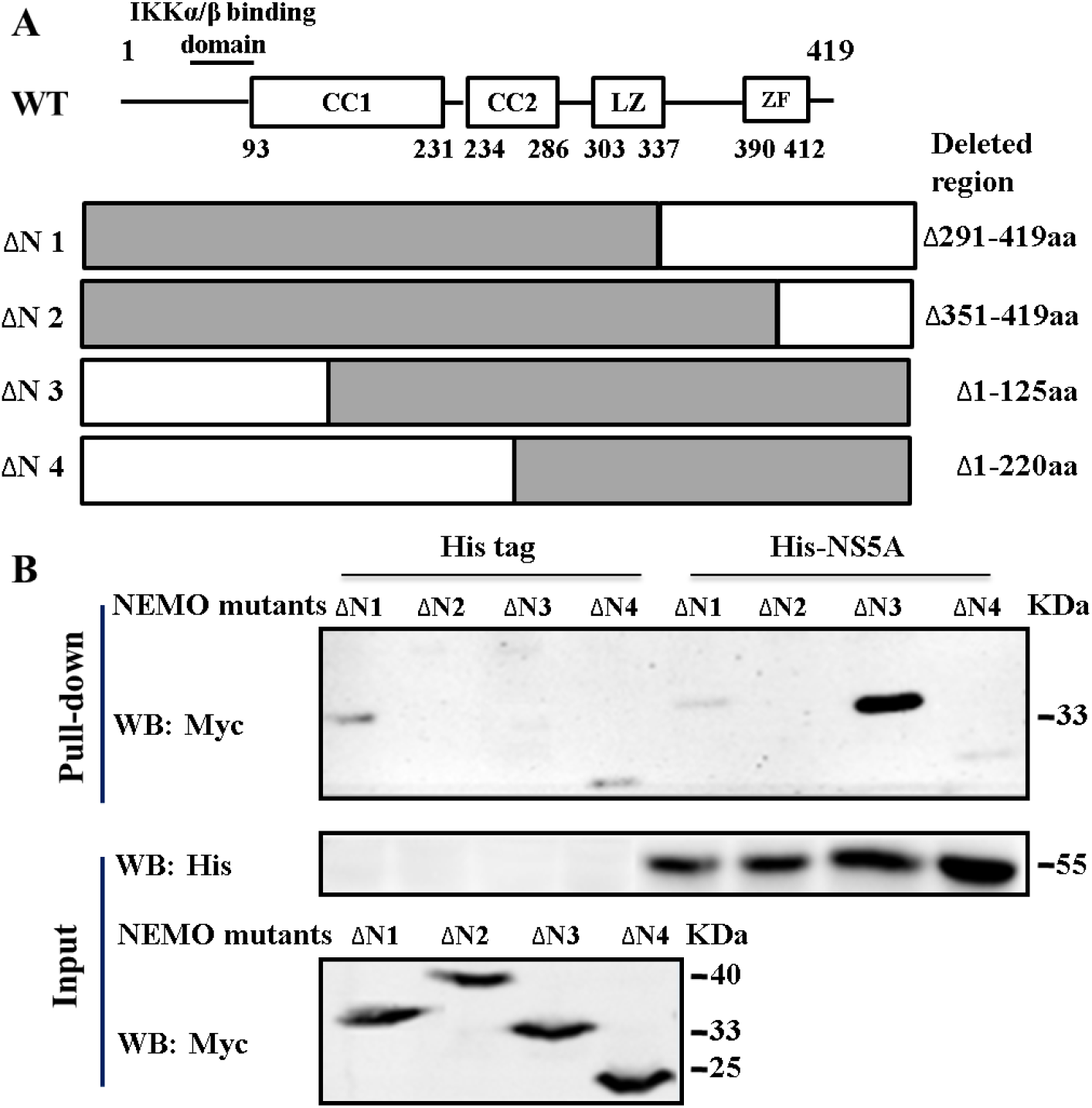
Determination of the domain of NEMO required for association with NS5A. (A) Schematic representation of NEMO mutant constructs. (B) 293T cells with expression of individual NEMO mutants were lysed and incubated with Dynabeads (Thermo Fisher Scientific, USA) coupled with His-NS5A or His tag. The proteins co-purified with fusion protein His-NS5A or His tag and cell lysates were subjected to western blot analysis using the indicated antibodies.

### The region of NS5A required for NS5A-NEMO interaction

293T cells were transfected with recombinant vectors expressing NS5A or one of 4 truncated mutants (p3xFlag-NS5Am1, p3xFlag-NS5Am2, p3xFlag-NS5Am3, p3xFlag-NS5Am4) (Fig 3A) or Flag tag and His-NEMO. At 48 h post-transfection, the cells were lysed, immunoprecipitated with mouse anti-His tag monoclonal antibody (Abcam, USA), and subjected to western blotting with rabbit anti-NEMO monoclonal antibody (Abcam, USA) and mouse anti-Flag monoclonal antibody (Proteintech, Wuhan, China). Fig 3B shows that only NS5Am2 (251-497 aa) failed to interact with NEMO, indicating that the 126-250aa segment of NS5A is responsible for its interaction with NEMO.

**Fig. 3.**
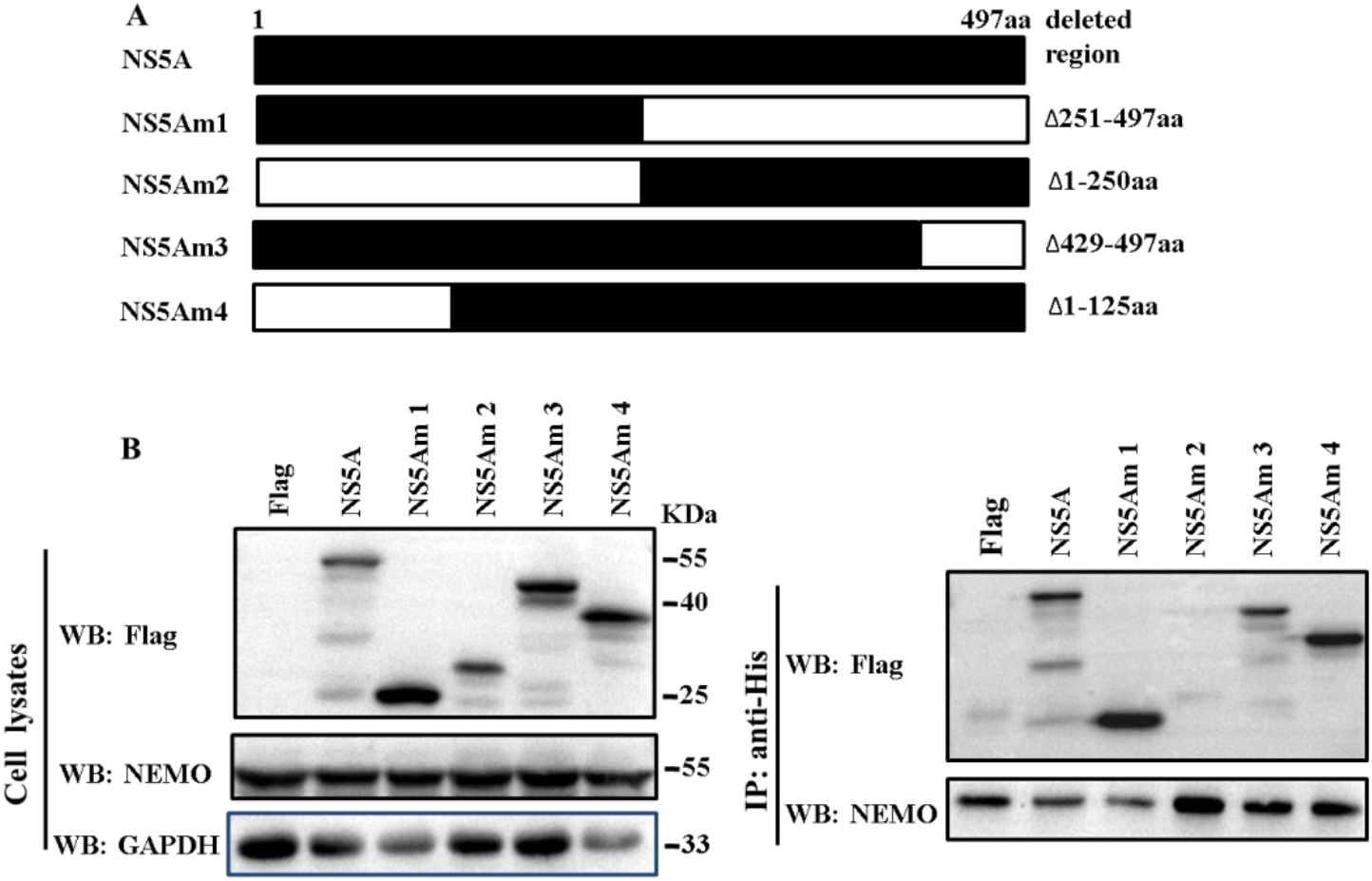
Determination of the region of NS5A required for NS5A-NEMO interaction. (A) Schematic representation of NS5A mutant constructs. (B) 293T cells expressing NS5A and an individual mutant or Flag tag were lysed for IP and western blot assay with the indicated antibodies.

### The region of NS5A required for inhibition of NF-κB signaling

293T cells were co-transfected with recombinant vectors expressing NS5A or individual truncated NS5A (p3xFlag-NS5Am1, p3xFlag-NS5Am2, p3xFlag-NS5Am3, p3xFlag-NS5Am4), pNFκB-Luc reporter and PRL-TK. At 48 h post-transfection, the cells were incubated with TNFα for 15 min, then harvested and assayed for NF-κB transcriptional activity. Fig. 4 A shows the expression of NS5A and the NS5A mutants with Fig. 4 B revealing that only the NS5Am2 mutant failed to inhibit TNFα-mediated NF-κB signaling.

**Fig. 4.**
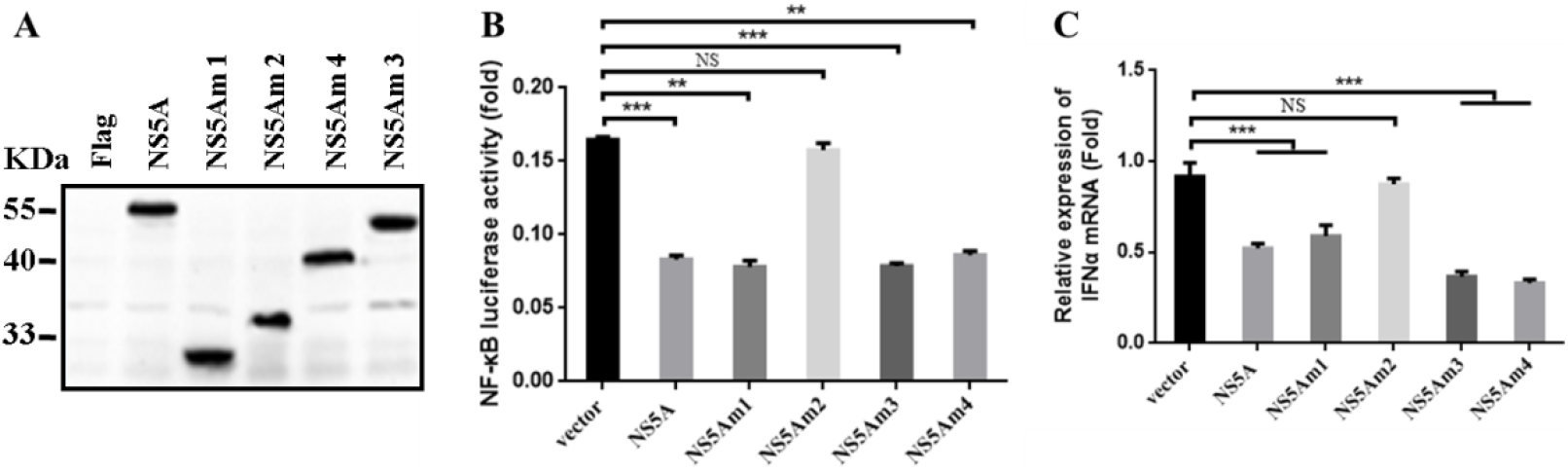
Determination of the region of NS5A required for inhibition of NF-κB signaling. (A) Western blot detection of NS5A and NS5A truncated mutants in 293T cells. (B, C) 293T cells were transfected with recombinant vectors expressing truncated NS5A mutants (p3xFlag-NS5Am1, p3xFlag-NS5Am2, p3xFlag-NS5Am3, p3xFlag-NS5Am4) (C) and pNFκB-Luc reporter and PRL-TK (B). At 48 h post-transfection, the cells were incubated with TNFα for 15 min, then harvested and assayed for NF-κB transcriptional activity (B) or quantification of IFNα mRNA (C).

Transfected 293T cells were used to evaluate the effect of NS5A or the NS5A truncated mutants on NF-κB dependent IFNα mRNA expression. At 48 h the transfected cells were incubated with TNFα for 15 mins, then harvested and assayed for IFNα mRNA expression. As before, only the NS5Am2 mutant failed to inhibit IFNα mRNA expression (Fig.4 B), indicating again that the 126-250 aa region was implicated in the inhibition of NF-κB signaling by NS5A.

### Reduction of NEMO induced by CSFV or NS5A

PK-15 cells overexpressing Myc-NEMO were infected with CSFV Shimen (virulent) strain at an MOI of 5 TCID_50_. At 48 hpi, the cells were incubated with TNFα for 15 min and then lysed for western blot analysis with rabbit anti-NEMO monoclonal antibody (Abcam, USA). Viral infection of the cells was detected by indirect immunofluorescence (Fig.5A). Fig.5B shows that CSFV infection reduces total NEMO protein in untreated or TNFα-treated cells with exogenous NEMO expression. Interestingly, a greater reduction of NEMO was observed in TNFα-treated cells.

**Fig. 5.**
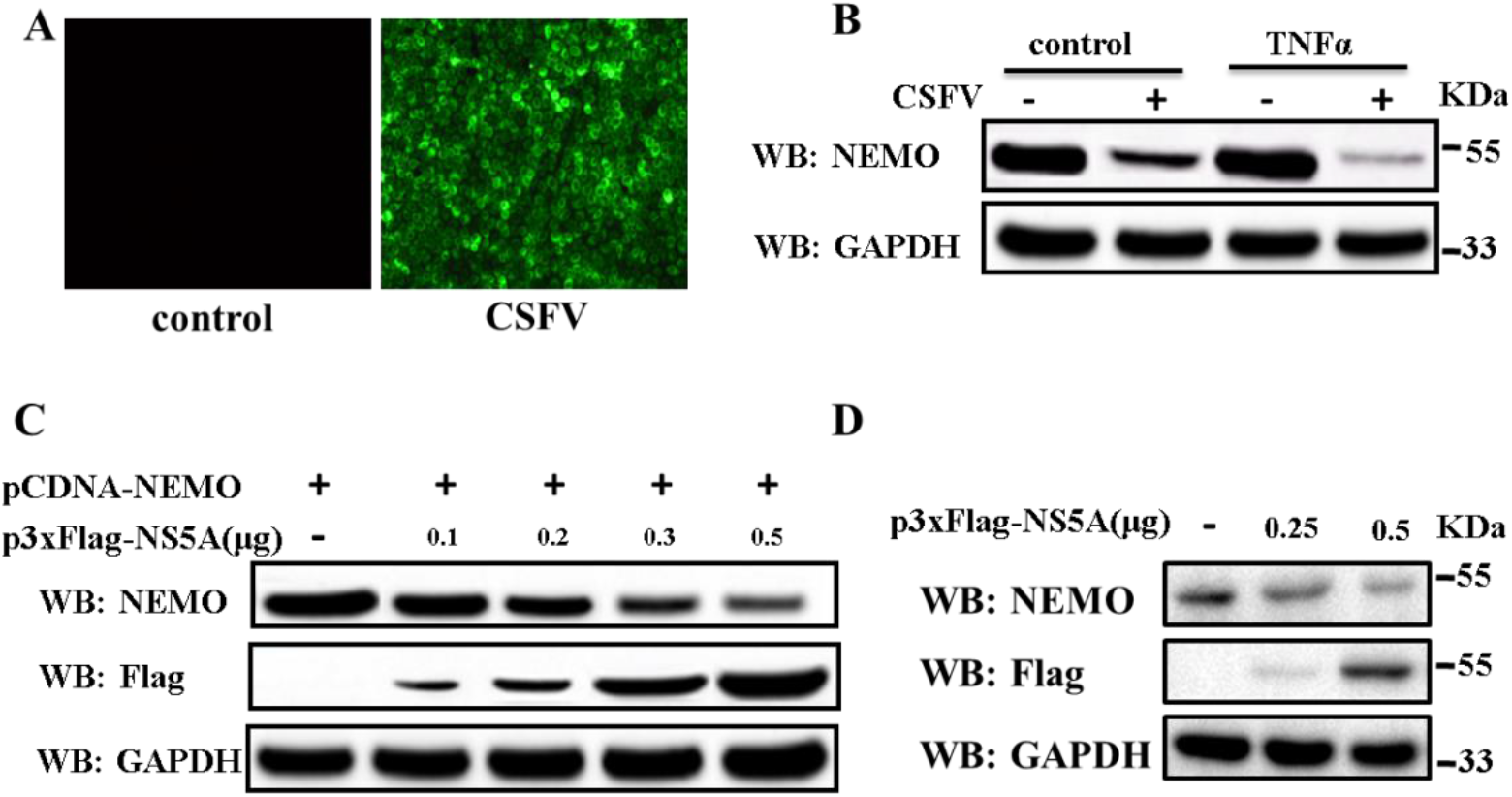
Inhibition of NEMO induced by CSFV or NS5A.(A)PK-15 cells were infected with CSFV Shimen strain at an MOI of 5 TCID_50_. At 48 hpi, CSFV infection was detected by indirect immunofluorescence. (B) PK-15 cells overexpressing Myc-NEMO were infected with CSFV Shimen strain at an MOI of 5 TCID_50_, at 48 hpi, the cells were incubated with TNFα or left untreated for 15 min before being lysed for western blot analysis with the indicated antibodies. (C) 293T cells were co-transfected with pCDNA-NEMO and increasing amounts of p3xFLAG-NS5A vectors. At 48 h post-transfection, the cells were incubated with TNFα for 15 min and lysed for western blot analysis with the indicated antibodies. (D) 293T cells were transfected with 0.25 and 0.5 µg of p3xFLAG-NS5A vectors. At 48 h post-transfection, the cells were treated with TNFα for 15 min and lysed for western blot analysis with the indicated antibodies.

293T cells were co-transfected with pCDNA-NEMO and increasing amounts of p3xFLAG-NS5A vectors. At 48 h post-transfection, the cells were lysed for western blot analysis with rabbit anti-NEMO monoclonal antibody (Abcam, USA). Fig.5C shows that levels of NEMO progressively decreased with increases in NS5A expression in cells with exogenous NEMO expression. Consequently, the effect of NS5A on endogenous NEMO was investigated, with results showing that NEMO progressively decreased as the expression of NS5A increased (Fig.5D).

### NS5A mediates proteasomal degradation of NEMO

The proteasome inhibitor MG132 was used to examine the possibility that NS5A mediates the proteasomal degradation of NEMO. As shown in Fig.6A, NEMO protein levels increased with increasing MG132 concentrations in cells with exogenous NEMO expression. Fig.6B also shows that MG132 inhibited endogenous NEMO degradation mediated by NS5A. These data indicate that NS5A induces proteasomal degradation of NEMO.

**Fig. 6.**
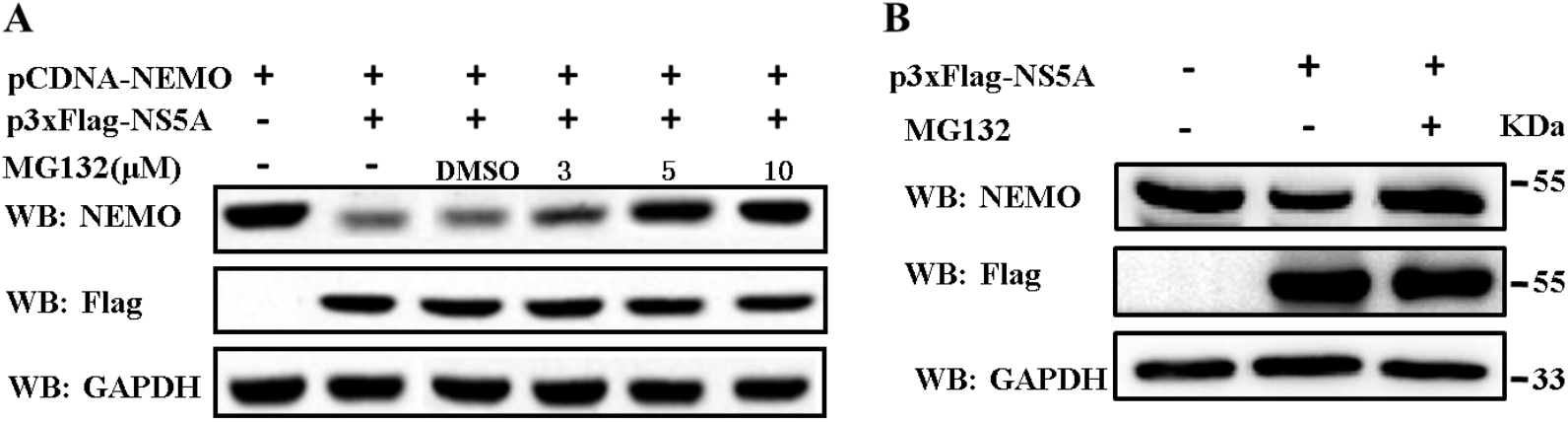
Evidence that NS5A mediates proteasomal degradation of NEMO. 293T cells were co-transfected with pCDNA-NEMO and p3xFLAG-NS5A vectors (A) or with p3xFLAG-NS5A vector alone (B). Following incubation with MG132 for 24 h and at 48 h post-transfection, the cells were lysed for western blot analysis with the indicated antibodies (A,B).

### NS5A mediates NEMO ubiquitination but does not mediate K48-linked polyubiquitination of NEMO

To test whether NS5A induces NEMO ubiquitination,293T cells were co-transfected with pCMV-HA-Ub and p3xFlag-NS5A or p3xFlag-CMV-10 (Fig.7A, B). At 48 h post-transfection, the cells were incubated with TNFα for 15 min and then lysed for immunoprecipitation with rabbit anti-NEMO monoclonal antibody (Abcam, USA). Precipitated proteins were subjected to western blotting using anti-HA antibody for probing ubiquitinated NEMO. Fig.7B shows that ubiquitinated NEMO increases in cells expressing NS5A, indicating that NS5A induces NEMO ubiquitination.

**Fig. 7.**
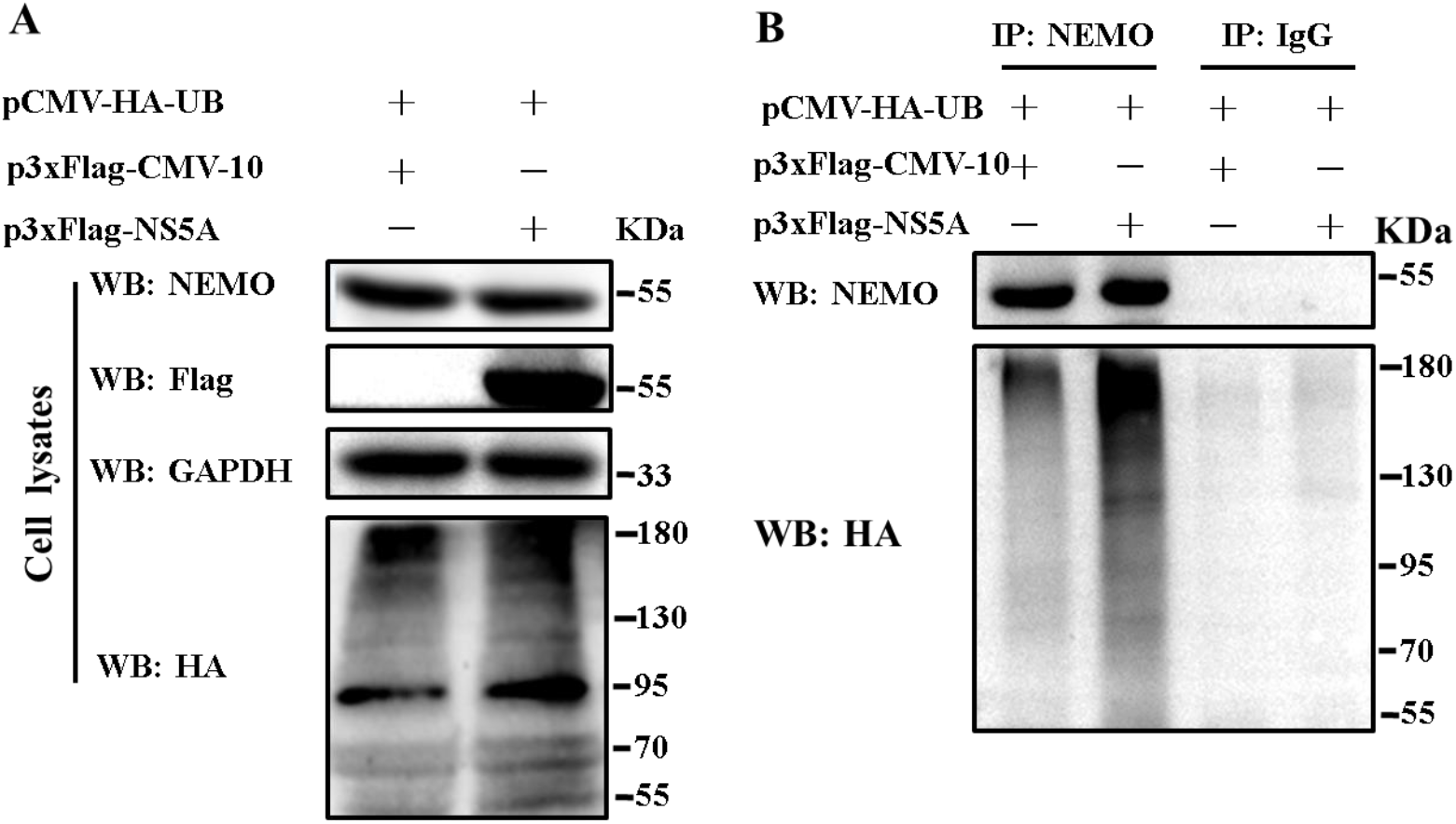
Evidence that NS5A mediates NEMO ubiquitination. 293T cells were transfected with the indicated vectors. At 48 h post-transfection, the cells were incubated with TNFα for 15 min and then lysed for immunoprecipitation (IP) with rabbit anti-NEMO monoclonal antibody or nonspecific IgG. Precipitated proteins and whole cell lysates were subjected to western blot analysis using the indicated antibodies.

K48-linked polyubiquitination usually triggers proteasomal degradation. To investigate whether NS5A mediates K48-linked polyubiquitination and subsequent degradation of NEMO, 293T cells were co-transfected with pCMV-HA-K48Ub and other indicated plasmid vectors (Fig.8). At 48 h post-transfection, the cells were incubated with TNFα for 15 min and then lysed for immunoprecipitation with mouse anti-His monoclonal antibody (Abcam, USA). Precipitated proteins were subjected to western blotting using anti-HA antibody for probing K48-linked polyubiquitinated NEMO (Fig.8). Results shows that there was no K48-linked polyubiquitinated NEMO in NS5A-expressed or control cells (Fig.8), indicating that NS5A does not mediate K48-linked polyubiquitination of NEMO.

**Fig. 8.**
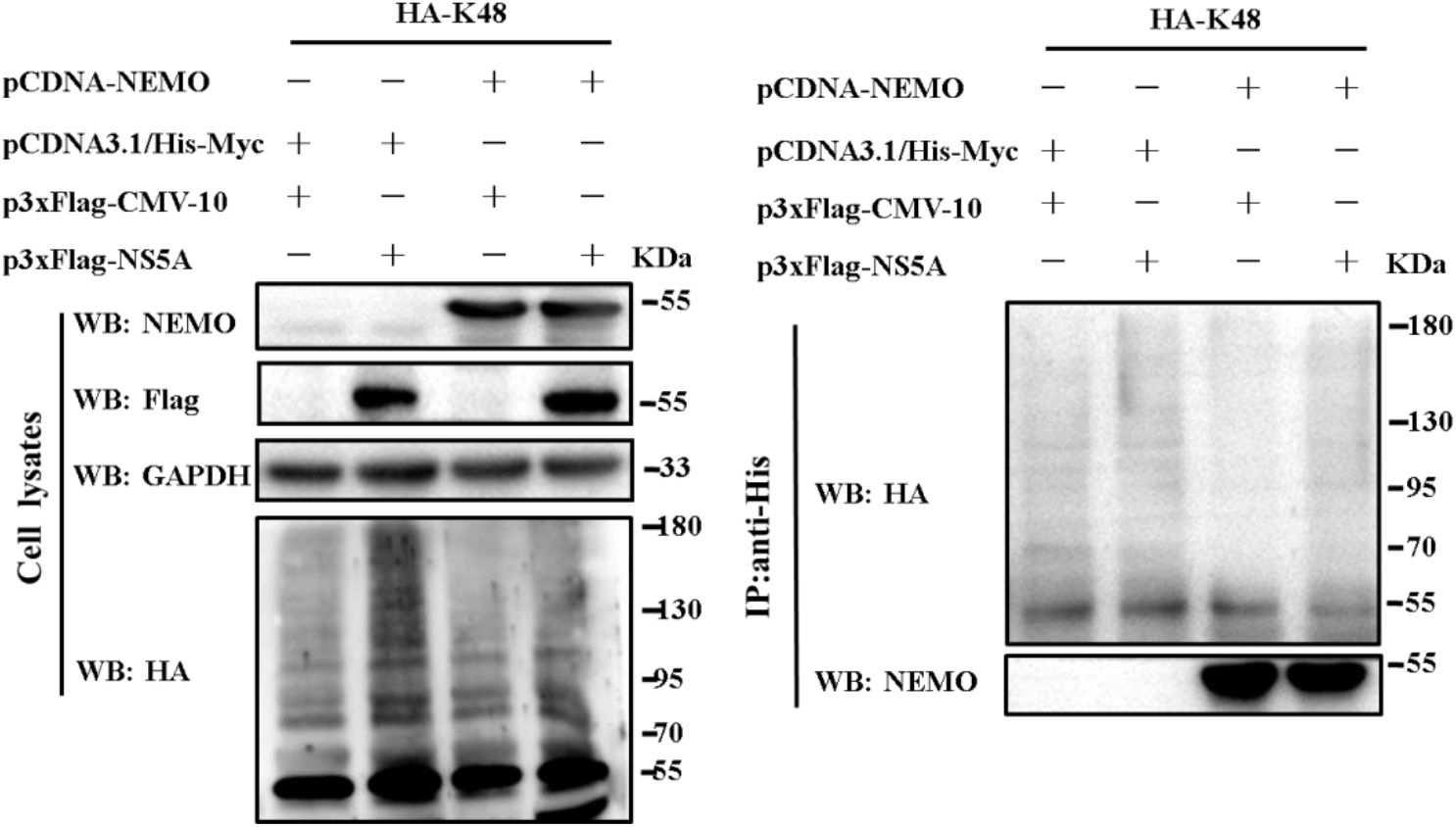
Evidence that NS5A does not mediate K48-linked polyubiquitination of NEMO. 293T cells were transfected with the indicated vectors. At 48 h post-transfection, the cells were treated with TNFα for 15 min and then lysed for immunoprecipitation (IP) with mouse anti-His tag monoclonal antibody. Precipitated proteins and whole cell lysates were subjected to western blot analysis using the indicated antibodies.

### NS5A induces K27-linked polyubiquitination of NEMO

To identify the type of polyubiquitination of NEMO for proteasomal degradation, ubiquitin (Ub) mutant genes (K6, K11, K27, K29, K33, and K63) were synthesized and cloned into the *Eco*RI and *Xho*I sites of pCMV-HA vector. Each mutant contained only one lysine (K) residue, with the 6 others having been replaced by arginine (R). Cultures of 293T cells were co-transfected with p3xFlag-NS5A, pCDNA-NEMO and pCMV-HA-Ub or individual Ub expression vector mutants (Fig.9A). At 48 h post-transfection, cells were incubated with TNFα for 15 mins followed by immunoprecipitation and ubiquitination assay of NEMO. Results shows a marked enhancement of polyubiquitination of NEMO in the presence of wild HA-Ub and mutant HA-Ub-K27 (Fig. 9A). To further validate that NS5A induced K27-linked polyubiquitination of NEMO, ubiquitin mutant HA-Ub-K27R was used. Fig. 9B shows that polyubiquitination of NEMO was eliminated in the presence of HA-Ub-K27R. These data indicat that NS5A induces K27-linked polyubiquitination of NEMO for proteasomal degradation.

**Fig. 9.**
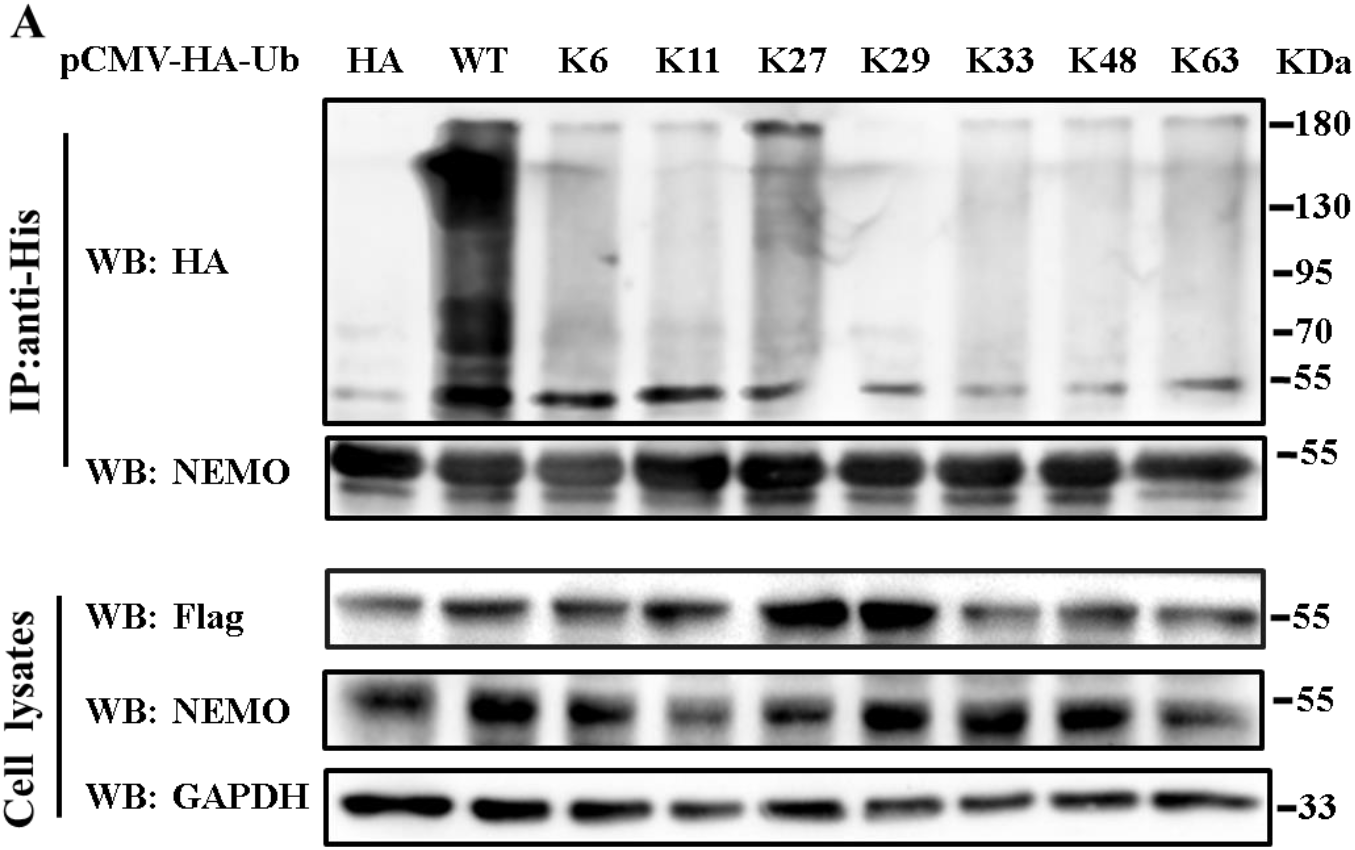
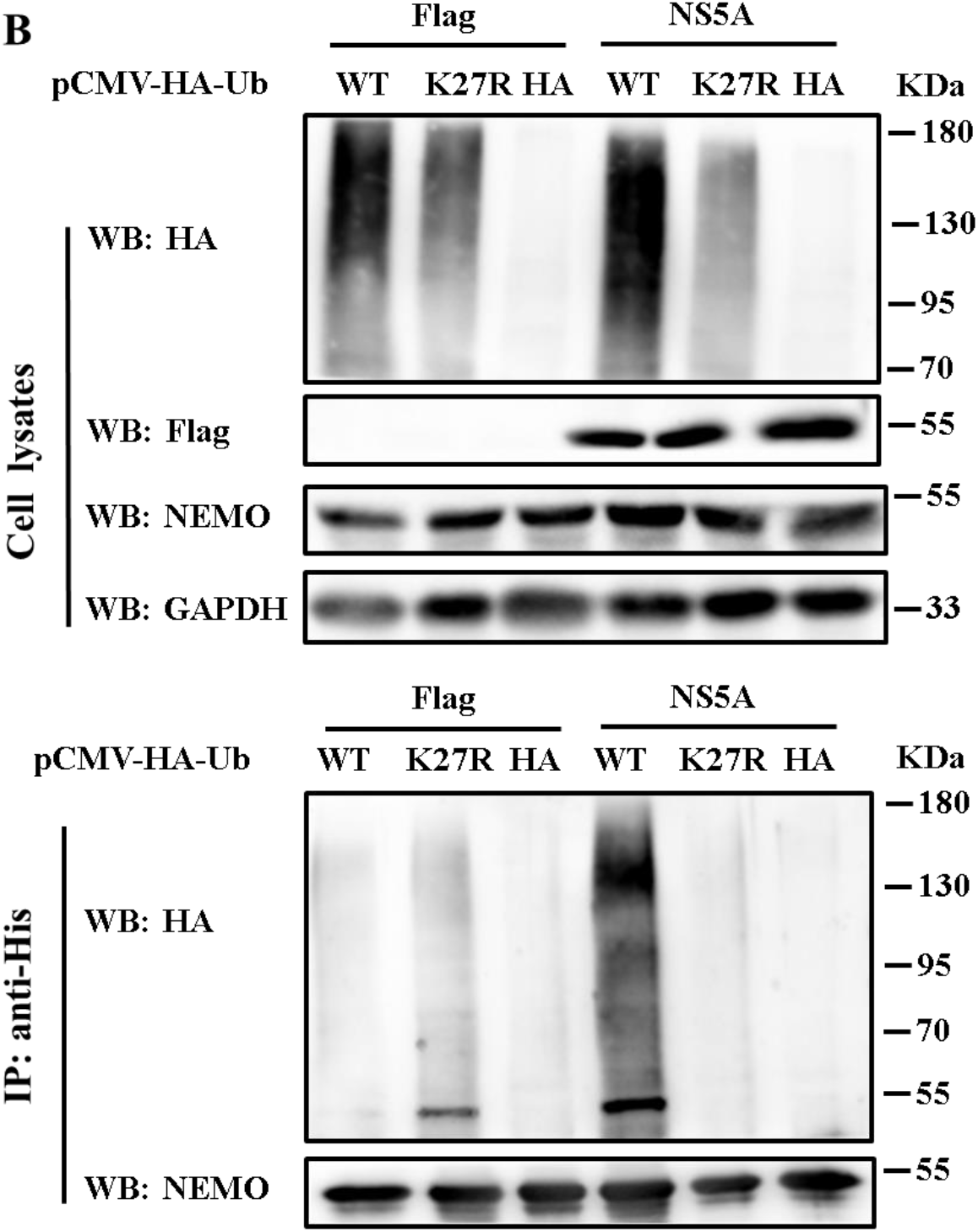
Evidence that NS5A induces K27-linked polyubiquitination of NEMO. (A) 293T cells were co-transfected with p3xFlag-NS5A, pCDNA-NEMO and pCMV-HA-Ub or individual Ub mutant expression vectors. At 48 h post-transfection, cells were treated with TNFα for 15 min before immunoprecipitation and ubiquitination assay of NEMO with the indicated antibodies. (B) 293T cells were co-transfected with p3xFlag-NS5A or p3xFlag-CMV-10, pCDNA-NEMO and pCMV-HA-Ub or pCMV-HA-K27R. At 48 h post-transfection, cells were incubated with TNFα for 15 min before immunoprecipitation and ubiquitination assay of NEMO with the indicated antibodies.

### NS5A inhibits K63-linked polyubiquitination of NEMO

As shown in Fig 2B, mapping the region of interaction of NEMO with NS5A had revealed that the NEMO C-terminal region of 351-419aa containing a zinc finger (ZF) domain is necessary for the interaction of NEMO with NS5A. The ZF domain of NEMO has been reported to be essential for ubiquitination of NEMO and IKK activation by TNFα (28, 29). To determine whether NS5A inhibits NEMO K63-linked polyunbiquitination, 293T cells were co-transfected with the indicated vectors (Fig.10A, B), incubated with TNFα for 15 min at 48 h post-transfection, then immunoprecipitated with anti-Myc antibody (Fig.10A) or rabbit anti-NEMO monoclonal antibody (Abcam, USA) and nonspecific IgG (Fig.10B). Precipitated proteins were subjected to western blotting with anti-HA antibody. Results showed that K63-linked polyunbiquition of exogenous NEMO (Fig.10A) and levels of endogenous NEMO (Fig.10B) were decreased significantly in NS5A-expressing cells.

**Fig. 10.**
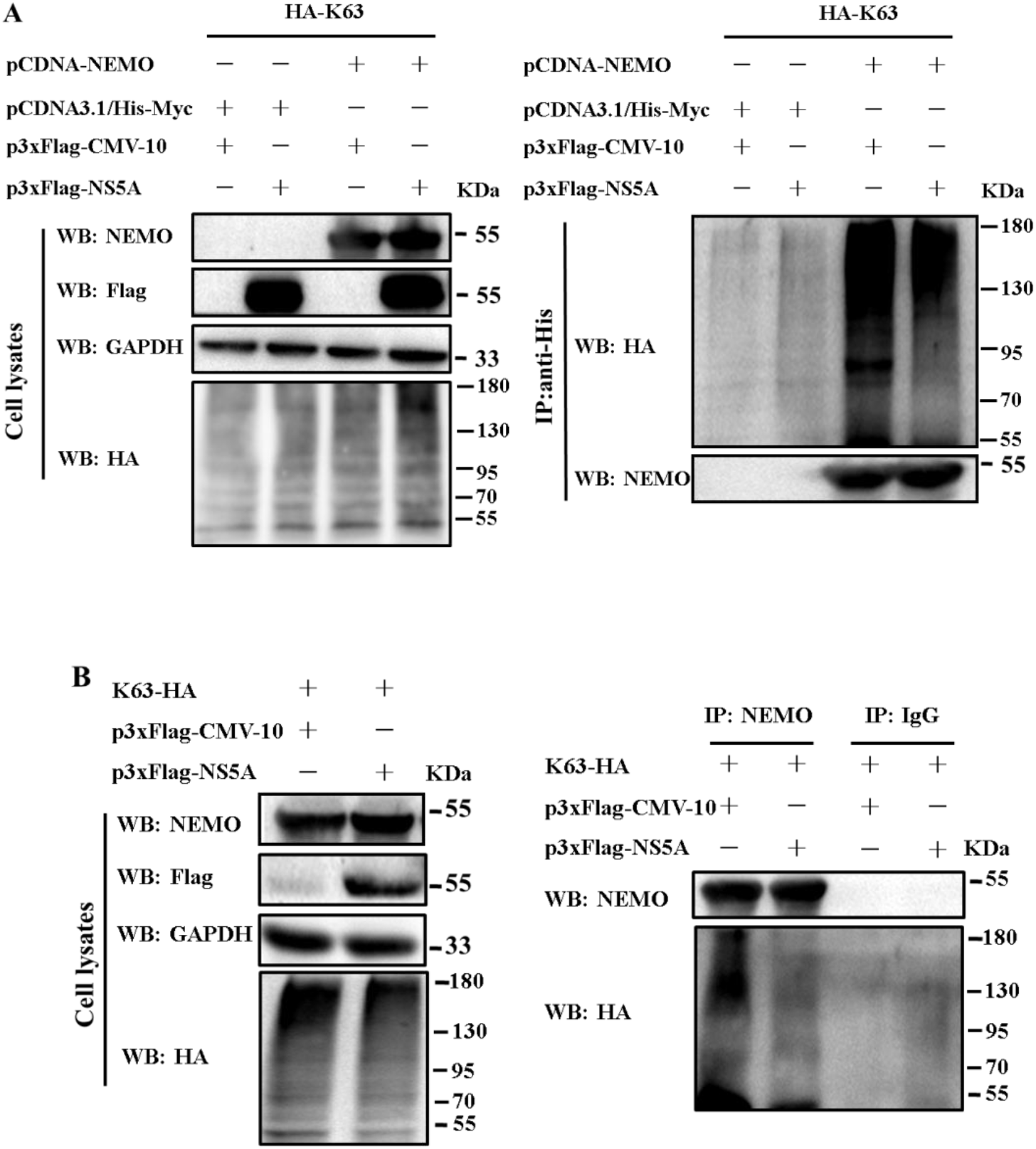
Evidence that NS5A inhibits K63-linked polyubiquitination of NEMO. 293 T cells were co-transfected with the indicated vectors (Fig. 10A, B). At 48 h post-transfection, cells were incubated with TNFα for 15 min before immunopreciptation with mouse anti-His monoclonal antibody (Fig. 10A) or anti-NEMO antibody and unspecific IgG (Fig. 10B). Precipitated proteins and whole cell lysates were subjected to western blot analysis using the indicated antibodiess.

## Discussion

NF-κB plays a critical role in inflammation and the innate immune responses (30). Many viruses therefore have developed various strategies targeting NF-κB to evade host immune response. Unexceptionally, CSFV infection fails to activate NF-κB signaling pathway (31) and NS5A protein suppresses NF-κB signaling induced by poly(I:C) (18). However, the mechanism of the inhibition of NF-κB signaling by NS5A is was not elucidated. Here, we reveal that NS5A mediates proteasomal degradation of NEMO and blocks k63-linked polyubiquitination of NEMO, thereby inhibiting NF-κB signaling.

Upon stimulation of canonical NF-κB signaling pathway by TNFα, NEMO mediates the binding of the IKK complex to K63-polyubiquitinated receptor-interacting protein 1(RIP1) by association with K63-linked polyubiquitin chains coupled to RIP1 for IKK phosphorylation by transforming growth factor-β-activated kinase-1 (TAK1) (20,21). Activated IKK subsequently phosphorylates IκBα, which triggers IκBα ubiquitination and subsequent proteasomal degradation, thereby allowing NF-κB to translocate to the nucleus and to trigger the transcription of its target genes. Dong & Tang (18) have reported that CSFV NS5A blocks degradation of IκBα and NF-κB nuclear translocation induced by poly(I:C), which indicates that NS5A probably targets an upstream event of IκBα phosphorylation, such as IKK complex formation, NEMO ubiquitination, or IKK phosphorylation, to inhibit NF-κB signaling.

NEMO’s N-terminal region binds to the C-terminal segment (NEMO-binding domain, NBD) of IKKα and IKKβ to form a IKK complex and modulates IKK complex activity (32). Disruption of the association between NEMO and IKK by a peptide, NBD mimetics or non-peptide small molecules inhibits the activation of IKK and NF-κB (33-35). Whether or not the interaction of CSFV NS5A with NEMO disrupts the association of NEMO with IKKα/IKKβ, interaction assays indicates that the N-terminal region (1-125 aa) of NEMO containing the IKKα/IKKβ binding domain (50–93 aa) (33) is unnecessary for its interaction with NS5A (Fig.2), which diminishes the likelihood that NS5A blocks assembly of the IKK complex by binding to NEMO.

The critical role of NEMO in the activation of NF-κB signaling makes it an attractive target for viruses to inhibit NF-κB signaling. It has been reported that some viruses, such as foot-and-mouth disease virus (36), porcine epidemic diarrhea virus (37) and porcine reproductive and respiratory syndrome virus (38) target NEMO and cleave it with their proteins with protease activity to inhibit the activation of IKK and NF-κB signaling. In addition, Brady et al. (39) have reported that molluscum contagiosum virus MC005 protein interacts with NEMO and blocks the conformational priming of the IKK kinase complex which occurs when NEMO couples to ubiquitin chains during NF-κB activation, thereby inhibiting IKK activation (39). Here, we have shown that levels of NEMO protein are reduced in CSFV-infected or NS5A-expressing cells (Fig.5), but recover in NS5A-expressing cells in the presence of proteasome inhibitor MG132 (Fig.6), indicating that NS5A mediated proteasomal degradation of NEMO.

K48-linked polyubiquitination being the major signal of proteasomal degradation, we therefore tested whether NS5A induced K48-linked polyubiquitination of NEMO. Ubiquitination assay demonstrated that NS5A induced K27-linked but not K48-linked polyubiquitination of NEMO for its proteasomal degradation (Fig.9). Further study is needed to clarify how NS5A induces K27-linked polyubiquitination of NEMO.

Ubiquitination of NEMO has been shown to be essential for the activation of IKK induced by TNFα (28, 40). Jun et al. (41) have shown that in addition to the binding of NEMO to polyubiquitin chains coupled to RIP1, site-specific ubiquitination of NEMO itself is essential for physiologic NF-kB signaling. Given the essential role of polyubiquitination of NEMO in the activation of IKK and NF-κB, viruses have evolved strategies to inhibit NF-κB signaling by blocking NEMO polyubiquitination. Biswas and Shisler (42) have reported that molluscum contagiosum virus (MCV) MC159 protein binds to the N-terminal region of NEMO and blocks the interaction of NEMO with cellular inhibitor of apoptosis protein 1 (cIAP1), a cellular E3 ligase mediating K63-linked polyubiquitination of NEMO, resulting in the inhibition of IKK and NF-κB activation. Wu et al. (43) have shown that the SARS-CoV-2 ORF9b N-terminus interacts with NEMO and blocks its K63-linked ubiquitination, leading to inhibition of NF-κB signaling and IFN production. Here, we have demonstrated that CSFV NS5A interacts with NEMO and that the C-terminal ZF domain of NEMO is required for the interaction (Fig. 2B). Significantly, the ZF domain of NEMO is a functional ubiquitin-binding domain (29) and an intact zinc finger of NEMO is essential for NEMO ubiquitination and the activation of IKK and NF-κB (28). Considering the important role of ZF in NEMO ubiquitination, the potential effects of NS5A on NEMO ubiquitination were analysed. Fig.10 shows that K63-linked polyubiquitination of NEMO is decreased in NS5A-expressing cells, indicating that NS5A inhibits K63-linked polyubiquitination of NEMO.

In summary, this study has identified CSFV NS5A as a novel antagonist of innate immunity and has shown that NS5A inhibits NF-κB signaling and type I IFN production by mediating proteasomal degradation of NEMO and blocking K63-linked polyubiquitination of NEMO. These findings reveal a novel mechanism by which CSFV evades host innate immunity and contribute to better understand of the pathogenesis of CSFV.

## METHODS

### Cell culture

Pig kidney cell line PK-15 and human kidney 293T cells were maintained in Dulbecco’s modified Eagle’s medium (DMEM) (Sigma, St. Louis, MO, USA) containing 10% fetal bovine serum (FBS; Gibco-BRL, Gaithersburg, MD, USA), 100 U penicillin G (Thermo Fisher Scientific, USA) and 100 µg streptomycin sulfate (Thermo Fisher Scientific, USA)/ml medium. The serum was negative for bovine viral diarrhea virus (BVDV) and BVDV antibody.

### Antibodies and reagents

Mouse anti-Flag monoclonal antibody, rabbit anti-Flag polyclonal antibody, mouse anti-GAPDH monoclonal antibody were purchased from Proteintech. Mouse anti-His tag monoclonal antibody, rabbit anti-NEMO monoclonal antibody were purchased from Abcam. Rabbit anti-NS5A serum was prepared in our laboratory. TNF-α, MG132 and *N*-ethylmaleimide were purchased from MedChemExpress. Lipo3000, Dynabeads-Protein G, Dynabeads™ His-Tag isolation and pulldown were purchased from Thermo Fisher Scientific.

### Plasmid construction

Construction of pET-NS5A and p3xFLAG-NS5A plasmids has been described previously (12). Four truncated mutants of NS5A (NS5Am1, NS5Am2, NS5Am3, NS5Am4) were amplified by PCR and inserted into the *Eco*RI and *Bam*HI sites of p3xFlag-CMV-10 vector to produce p3xFlag-NS5Am1, p3xFlag-NS5Am2, p3xFlag-NS5Am3, p3xFlag-NS5Am4. For overexpression of NEMO,its gene was synthesized and cloned into the *Eco*RI and *Xho*I sites of pcDNA3.1/*myc*-HisA vector (Invitrogen, Carlsbad, CA, USA) to produce pcDNA-NEMO. Four truncated mutants of NEMO (ΔN1, ΔN2, ΔN3, ΔN4) were amplified by PCR and inserted into the *Eco*RI and *Kpn*I sites of pCMV-Myc vector to produce pCMV-NEMOΔN1, pCMV-NEMOΔN2, pCMV-NEMOΔN3, pCMV-NEMOΔN4. Wild type ubiquitin gene and ubiquitin mutants genes (K6, K11, K27, K29, K33, K63) were synthesized and were inserted into the *Eco*RI and *Xho*I sites of pCMV-HA vector to produce pCMV-HA-Ub, pCMV-HA-K6Ub, pCMV-HA-K11Ub, pCMV-HA-K27Ub, pCMV-HA-K29Ub, pCMV-HA-K33Ub, and pCMV-HA-K63Ub. UbK27R gene was amplified by overlapping PCR and was in inserted into the *Eco*RI and *Xho*I sites of pCMV-HA vector to produce pCMV-HA-UbK27R. Primers used in present study were listed in table 1.

**Table 1.**
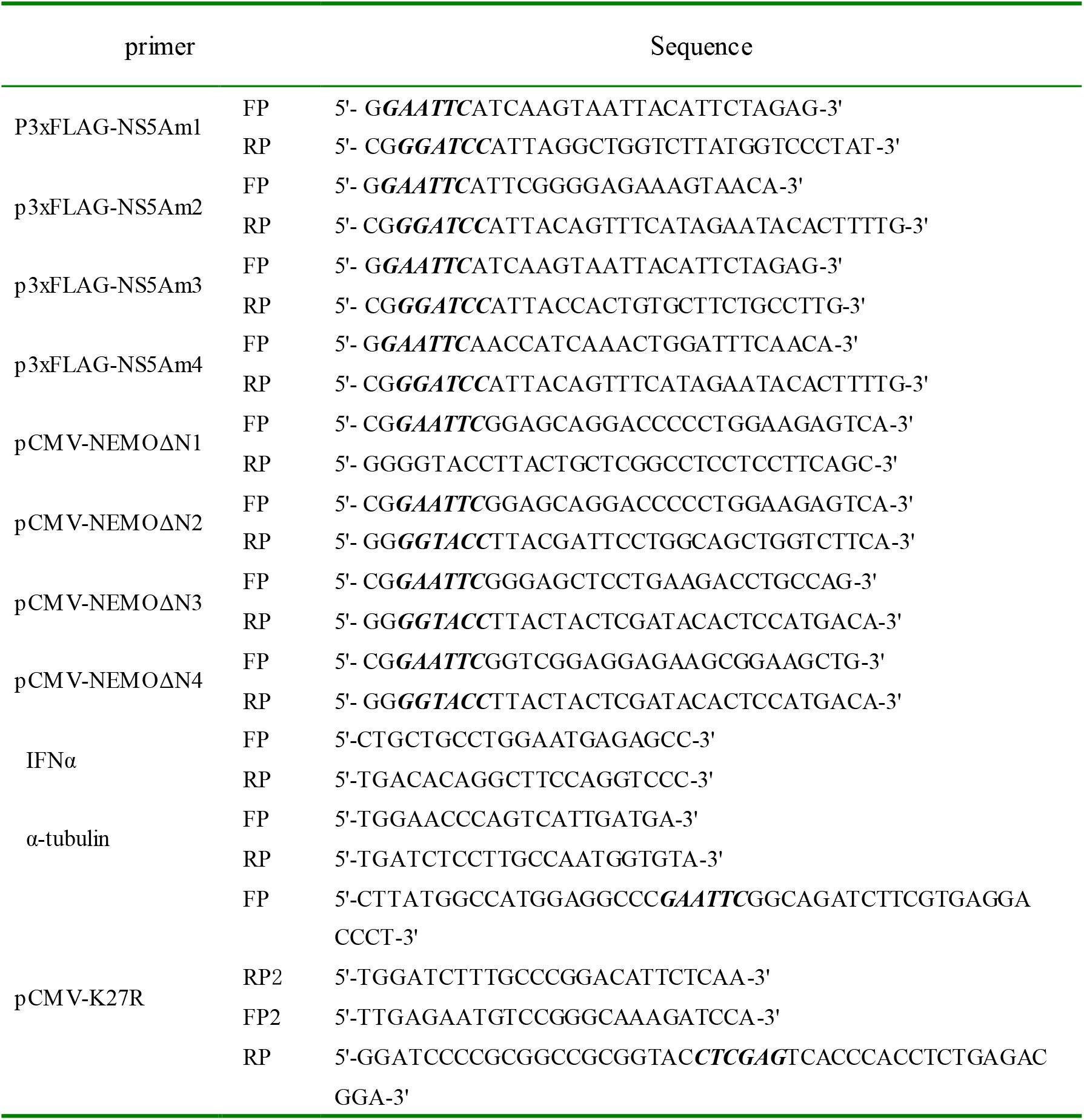
Sequences of oligonucleotide primers used in present study.

### Coimmunopreciptation (Co-IP)

Cells were transfected with recombinant plasmid p3xFLAG-NS5A or NS5A mutant expression vectors using Lipo3000 (Thermo Fisher Scientific, USA) according to the manufacturer’s directions. At 48 h post-transfection, total cell protein was extracted and subjected to co-immunopreciptation (Co-IP) with Dynabeads-Protein G (Thermo Fisher Scientific, USA) followed by the protocol described previously (12).

### His tag pull down assay

Pull down was conducted as described previously (12). Briefly, Dynabeads (Thermo Fisher Scientific, USA) with bound His-NS5A or His tag protein were incubated for 10 min on a roller at ambient temperature with cell lysates from 293T cells expressing Myc-NEMO. Proteins co-purified with His-NS5A or His tag protein were detected by western blot with mouse anti-Myc tag monoclonal antibody (Abcam, USA) for probing NEMO and mouse anti-His tag monoclonal antibody (Abcam, USA) for probing NS5A.

### Western blotting

Protein concentrations were determined with the BioTeke Protein Assay kit (Bioteke Corporation, Beijing, China) and subjected to western blotting following the previous protocol (Sun et al., 2008).

### NF-κB luciferase reporter assay

293T cells in 24-well plates were transfected with p3xFLAG -NS5A or p3xFLAG-CMV-10 vector at 1 µg per well along with pNFκB-luc (Beyotime Biotechnology, China) at 0.5 µg and pRL-TK (Beyotime) at 0.25µg per well. At 48 h post-transfection, the cells were stimulated with TNFα (final concentration 20 ng·mL^-1^) for 15 min and then harvested for firefly luciferase (FLuc) and Renilla luciferase (RLuc) activities assay using the Dual-Lumi™ Luciferase Reporter Gene Assay Kit (Beyotime).

### Real-time RT-PCR

293T cells in 24-well plates were transfected with p3×FLAG-NS5A or p3 ×FLAG-CMV-10 vector at 1 μg per well.At 48 h post transfection,the cells were simulated with TNF α (final concentration 20 ngmL ^-1^) for 15 min and then harvested for quantification of INF α mRNA using the One-step SYBR Prime script RT-PCR kit (TaKaRa, Dalian, China) according to the manufacturer’s protocol.Briefly the total RNA was isolated from TNF-α-treated with Trizol reagent (Takara Bio, Daian, China). cDNA was synthesized from 1 μg of total RNA with Prime Script II (Takara Bio, Daian, China). Quantitative real-time PCR analysis was performed using an ABI PRISM 7000 cycler (Applied Biosystems, Carlsbad, CA, USA) using FastStart Universal SYBR Green Master (Takara). The level of β-actin expression was used to normalize the data. The primers were listed in table 1.

### Ubiquitination assy

293T cells were transfected with p3xFLAG -NS5A or p3xFLAG-CMV-10, pCDNA-NEMO or pCDNA3.1/His-Myc and vectors expressing the different ubiquitin constructs and incubated with MG132 (MedChemExpress, USA) for 24 h. At 48 h post-transfection, the cells were stimulated with TNFα (final concentration 20 ng·mL^-1^) for 15 min and then lysed with lysis buffer containing 10 µM *N*-ethylmaleimide (MedChemExpress, USA), a inhibitor of deubiquitinating enzyme, for immunoprecipitation with mouse anti-His tag monoclonal antibody (Abcam, USA) or rabbit anti-NEMO monoclonal antibody (Abcam, USA). Precipitated NEMO was detected by western blot with rabbit anti-HA polyclonal antibody (Proteintech, China) for probing ubiqitination of NEMO.

### Statistical analysis

All experiments were conducted with at least three independent replicates. Experimental data were analyzed by GraphPad Prism software using Student’s *t* test. Differences in data were regarded as significant if P<0.05.

## Acknowledgements

This work was supported by the National Nature Science Fundation of China (Grant No.31972677).

